# mlplasmids: a user-friendly tool to predict plasmid- and chromosome-derived sequences for single species

**DOI:** 10.1101/329045

**Authors:** Sergio Arredondo-Alonso, Malbert R. C. Rogers, Johanna C. Braat, T. D. Verschuuren, Janetta Top, Jukka Corander, Rob J.L. Willems, Anita C. Schürch

## Abstract

Assembly of bacterial short-read whole genome sequencing (WGS) data frequently results in hundreds of contigs for which the origin, plasmid or chromosome, is unclear. Long-read sequencing has emerged as a solution to resolve plasmid structures and to obtain complete genomes for most bacterial species. This information can be used to generate and label datasets from short-read based contigs as plasmid- or chromosome-derived. We investigated the use of several popular machine learning methods to classify short-read contigs with known plasmid- or chromosome-origin from *Enterococcus faecium, Klebsiella pneumoniae* and *Escherichia coli* using pentamer frequencies. Based on resulting F1-scores we selected support-vector machine (SVM) models as best classifier for all three bacterial species (F1-score *E. faecium* = 0.94, F1-score *K. pneumoniae* = 0.90, F1-score *E. coli* = 0.76), which outperformed other existing plasmid tools using an independent set of isolates (precision *E. faecium* = 0.92, precision *K. pneumoniae* = 0.86, precision *E. coli* = 0.82). We demonstrated the scalability of our model by accurately predicting the plasmidome of a large collection of 1,644 *E. faecium* isolates with only short-read WGS available using a standard laptop with a single core. A low number of false positive predicted sequences suggests that the assignment of a particular gene of interest as plasmid- or chromosome-encoded by the models is plausible. The SVM classifiers are publicly available as a new R package called ‘mlplasmids’ at https://gitlab.com/sirarredondo/mlplasmids under the GNU General Public License v3.0. We additionally developed a graphical-user interface using the Shiny package which can be accessed at https://sarredondo.shinyapps.io/mlplasmids/. Single genomes can easily be predicted by uploading genome assemblies. We anticipate that this tool may significantly facilitate research on the dissemination of plasmids encoding antibiotic resistance and/or contributing to host adaptation.

## Introduction

Plasmids are autonomous extra-chromosomal elements which can act as major drivers of variation and adaptation in bacterial populations (1, 2). Plasmids can also facilitate the dissemination of antimicrobial resistance (AMR) via horizontal transfer of plasmids encoding resistance genes such as plasmid-derived vancomycin resistance in *Enterococcus faecium* or extended-spectrum beta-lactamase in *Enterobacteriaceae* isolates (3–6). This means that understanding plasmid epidemiology is pivotal to fully understand the introduction and transmission of AMR in bacterial populations (7, 8).

Analyzing the plasmid content of large collections of isolates by PCR-based techniques is laborious and has low resolution. Illumina sequencing platforms, which provide short reads (ranging from 150-300 bp) with low error rates, have been massively used to assemble bacterial draft genomes (9). However, the frequent presence of insertion-sequences (IS) and transposable elements in bacterial genomes prohibit their full assembly because these regions cannot be spanned by short-reads (7, 10). This results in a fragmented assembly typically consisting of hundreds of chromosomal and plasmid contigs that challenge the inference of the origin of these contigs.

Different tools (PlasmidFinder, cBAR, Recycler, PlasmidSPAdes, PlasFlow) have been proposed to automate the reconstruction of plasmids using short-read WGS data (11–15). However, plasmid predictions are usually incomplete and chromosome-derived contigs are frequently present among the predicted plasmids (16). This may be partially overcome using tools such as PlacnetW, which allows users to define and solve plasmid boundaries but limits the high-throughput analysis of short-read WGS data (17, 18).

Long-read WGS has emerged as a solution to obtain complete and error-free plasmid sequences (19, 20). Read lengths obtained by these platforms allow to completely span repeat sequences and obtaining a single contig per replicon (21, 22). Due to the increasing number of complete genomes available in RefSeq/NCBI databases, we explored the possibility of training several popular machine learning algorithms using genome signatures from single-species assemblies. These features have been previously used in cBAR and recently in PlasFlow to distinguish plasmid- and chromosome-derived sequences in metagenomic samples.

Here, we present mlplasmids, a new tool to predict plasmid- and chromosome-derived sequences for a selection of Gram-positive and Gram-negative bacterial species (*Enterococcus faecium, Klebsiella pneumoniae and Escherichia coli*) with species-specific classifiers and we show that mlplasmids outperforms other plasmid prediction tools for these three species. We made the plasmid models available as an R package and a web-server.

## Materials and methods

### Retrieving complete genome sequences from NCBI database

We downloaded complete genomes for *E. faecium* (chromosomes = 24; plasmids = 82), *K. pneumoniae* (chromosomes = 156; plasmids = 561) and *E. coli* (chromosomes = 168; plasmids = 415) from Assembly Entrez NCBI database (https://www.ncbi.nlm.nih.gov/assembly/) with the following criteria: i) an status level of ‘Complete Genome’ and ii) one or more plasmid entries in its respective genome assembly. Retrieved genomes and their corresponding accession sequences are available in Supplementary File 1.

### Extending the number of complete genome sequences for *E. faecium*

From a collection of 1,644 *E. faecium* Illumina-sequenced (MiSeq/NextSeq) isolates, we used Oxford Nanopore Technologies (ONT) to obtain complete genome sequences of 60 isolates. These isolates were selected based on their preliminary plasmid content using PlasmidSPAdes (version 3.8.2) and presence of known plasmid replication genes (23) (Supplementary Methods 1). Hybrid assembly using Unicycler (version 0.4.1) specifying ‘bold’ mode was used to obtain complete genome sequences derived from Illumina NextSeq and ONT reads. *Unicycler* contigs were categorized either as chromosome or plasmids based on size. Contigs were categorized as chromosome if they were larger than 350 kbp. Contigs smaller than 350 kbp were categorized as plasmids if they were circular. If contigs were smaller than 350 kbp but were lacking a circularization signal, they were not categorized. Detailed explanation about Illumina and ONT sequencing is available at Supplementary Methods 1.

### Estimating strain diversity in our collection of complete genomes

We estimated the diversity present in our collection of *K. pneumoniae, E.coli* and *E. faecium* genomes with Mash (version 1.1) (sketch size = 1,000; k-mer = 21) (24). For each isolate present in the collections of *E. faecium, K. pneumoniae* and *E. coli,* we computed pairwise Mash distances of all isolates belonging to a particular collection. Mash distances were calculated considering the total genome content of an isolate (chromosome plus associated plasmids). Computed pairwise Mash distances were transformed into a distance matrix and clustered using hclust function (method = ‘ward.D2’) available in R package stats (version 3.3.3). Hierarchical clustering was visualized using heatmap.2 function available in R package gplots (version 3.0.1) (25).

### Simulating Illumina sequence reads

For each selected complete genome of *K. pneumoniae* and *E. coli*, we retrieved its genome size using bioawk (version 20110810) to calculate the number of paired reads required to simulate sequence read files using wgsim (version 0.3.2) (https://github.com/lh3/wgsim) with 50x coverage and no error rate.

### Assembling Illumina sequence reads

Simulated sequence reads of *K. pneumoniae* and *E. coli*, were trimmed using seqtk (version 1.2-r94, https://github.com/lh3/seqtk) with the command “--trimfq”. We used SPAdes (version 3.6.2) to perform *de novo* assembly. Contigs with a length smaller than 500bp were excluded

*E. faecium* Illumina NextSeq reads were trimmed using *nesoni clip*, part of the *nesoni* toolkit (version 0.132), with the following settings: ‘--adaptor-clip yes --match 10 --max-errors 1 --clip-ambiguous yes --quality 10 --length 90 --trim-start 0 --trim-end 0 --gzip no --out-separate yes pairs:’. Trimmed reads were then assembled into contigs using SPAdes (version 3.5.0) with default settings. Contigs with an average coverage lower than 10x and/or a length smaller than 500bp were removed from the assemblies.

### Labeling short-read contigs as chromosome- or plasmid-derived

To label contigs as either plasmid- or chromosome-derived, SPAdes contigs were mapped using bwa-mem (version 0.7.15-r1140) against complete chromosomal and plasmid sequences. Contig alignments were parsed using samtools (version 1.4). This approach allowed to label each SPAdes contig either as plasmid- or chromosome-derived. SPAdes contigs mapping both to complete chromosomal and plasmid sequences or with a length shorter than 500 bp were discarded.

### Defining an independent set of isolates for validation

To create an independent set of isolates for validation of mlplasmids, we excluded isolates of *K. pneumoniae* (chromosomes = 10; plasmids = 33) and *E. coli* (chromosomes = 3; plasmids = 7) in which original Illumina sequencing data were publicly available and that we previously used to benchmark plasmid prediction tools (16) (Supplementary File 2). For the *E. coli* dataset we included SPAdes contigs from ‘*Escherichia coli* str. K-12 substr. MG1655’ to observe the performance of our *E. coli* classifier in a strain bearing no plasmids in the validation set. SPAdes genome assemblies were used to evaluate the performance of resulting *K. pneumoniae* and *E. coli* models in contigs resulting from non-simulated sequence reads.

For *E. faecium*, labeled SPAdes contigs derived from 5 isolates were excluded from the training set. This included negative control isolates where no plasmids were assembled after hybrid assembly (E2079, E2364, E9101) and two isolates containing plasmid sequences (E7591, E8172) which included the isolate bearing the highest number of plasmids (n = 11). We also included two other strains in the independent validation set that are part of the collection of *E. faecium* strains with available ONT reads (E4239, E4457). Publicly available complete *E. faecium* genomes (chromosomes = 24; plasmids = 82) were not considered to train the machine-learning models and were used to observe the performance of the machine-learning classifier in sequences with a length larger than average short-read contig.

### Genomic signatures as features to distinguish plasmid and chromosome sequences

To investigate the role of k-mers as classifier features to differentiate between plasmid and chromosomal sequences, we computed Mash distances (sketch size = 400; k-mer = 16) from all chromosomal and plasmid sequences present in the collections *E. coli*, *K. pneumoniae* and *E. coli* genomes. We used the same parameters as used in the original Mash publication to cluster RefSeq complete genomes (24). Computed pairwise Mash distances between all the sequences were transformed into a distance matrix and clustered using hclust function (method = ‘ward.D2’). Hierarchical clustering was visualized using heatmap.2 function available in R package gplots. Additionally, we used the t-Distributed Stochastic Neighbor Embedding (t-SNE) (theta = 0.5, iterations = 1000, dims = 2, is_distance = TRUE) using the implementation provided in the R package Rtsne (version 0.13) to cluster and visualize chromosomal and plasmid sequences using computed Mash distances (26).

### Building a machine-learning model

For each bacterial species, we tuned and compared five different supervised algorithms provided in mlr R package (version 2.11): logistic regression, Bayesian classifier, decision trees, random forest (RF) and support-vector machine (SVM) (27). We defined a two-class classification problem using the category ‘plasmid’ as positive-class. To train and test the resulting classifiers we considered pentamer frequencies (n=1024) which were calculated using oligonucleotideFrequency function available in R package biostrings (version 2.42.1). Mlr package was used to split SPAdes labeled contigs into training (80%) and test set (20%), preserving the frequencies of each class in both sets (Table 1).

**Table 1.**
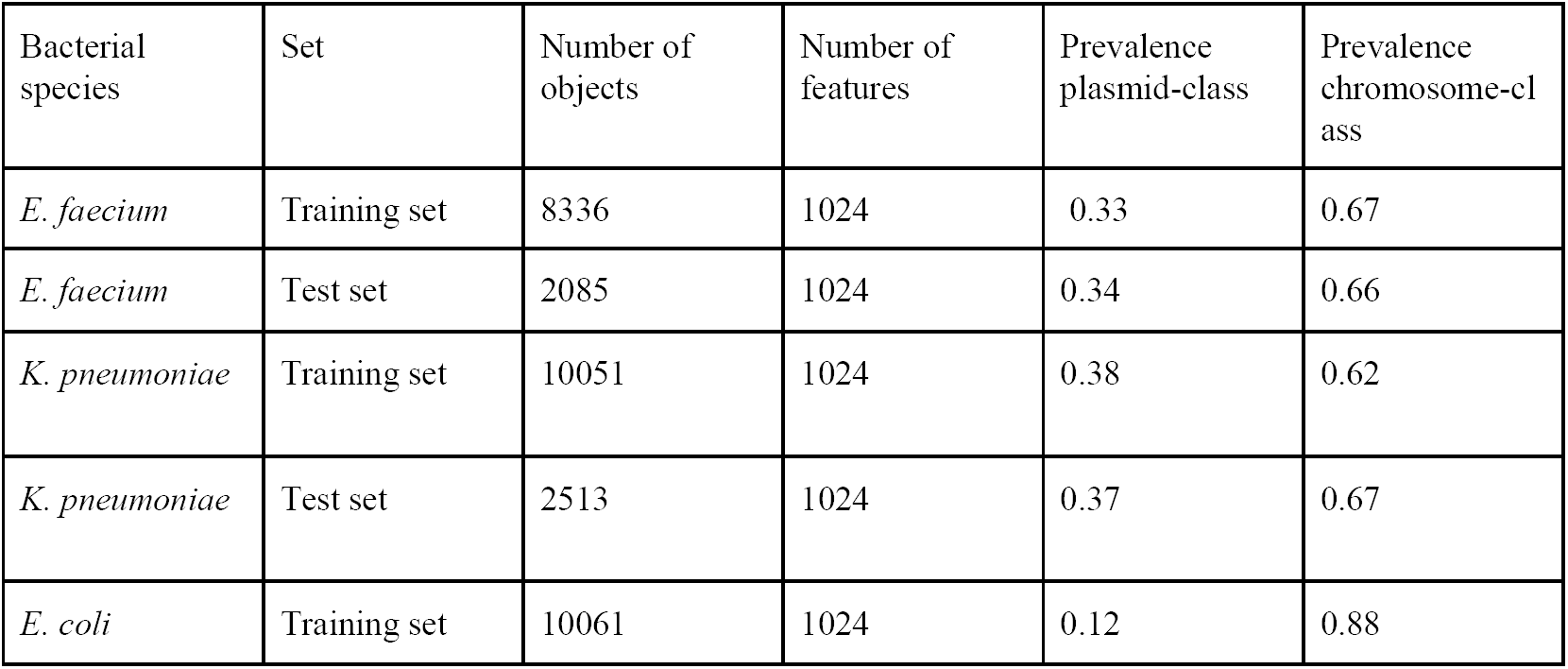
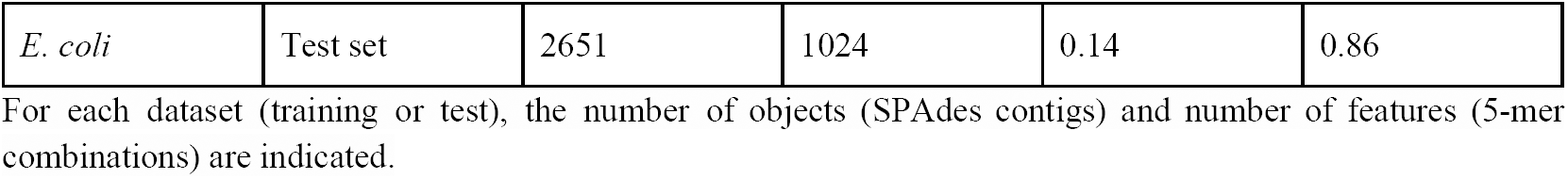
Description of the training and test sets used for each bacterial species.

Decision trees, random forest and support-vector machines hyperparameters were optimized using random search in a predefined search space (Supplementary Table S2). We performed 10-fold cross-validation to assess the quality of hyperparameters combination, using error rate as performance measure, except for *E. coli* models in which true positive rate was considered to overcome a lower plasmid frequency. For each object, posterior probabilities were generated and the class with a highest posterior probability was assigned.

Optimized classifiers were compared for the test set through receiver operating characteristic (ROC) curves. For each classifier, area under the curve (AUC) and precision-recall curves were calculated to compare resulting classifiers based on different true positive and false positive thresholds (from 0 to 1). Metrics were assessed using two different units: number of contigs and sequence size in base-pairs. F1 score was reported to obtain a harmonic average between specificity and sensitivity. Definitions of the statistics reported in this study are reported below.

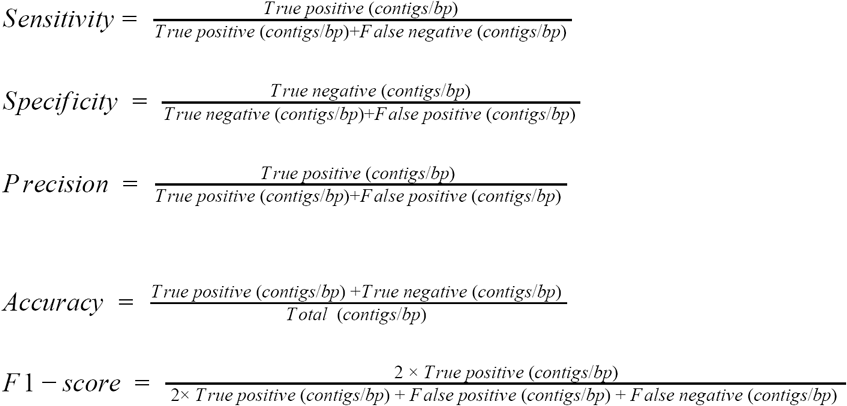

An overview of the method followed to build the resulting classifiers is shown in Figure 1.

**Figure 1.**
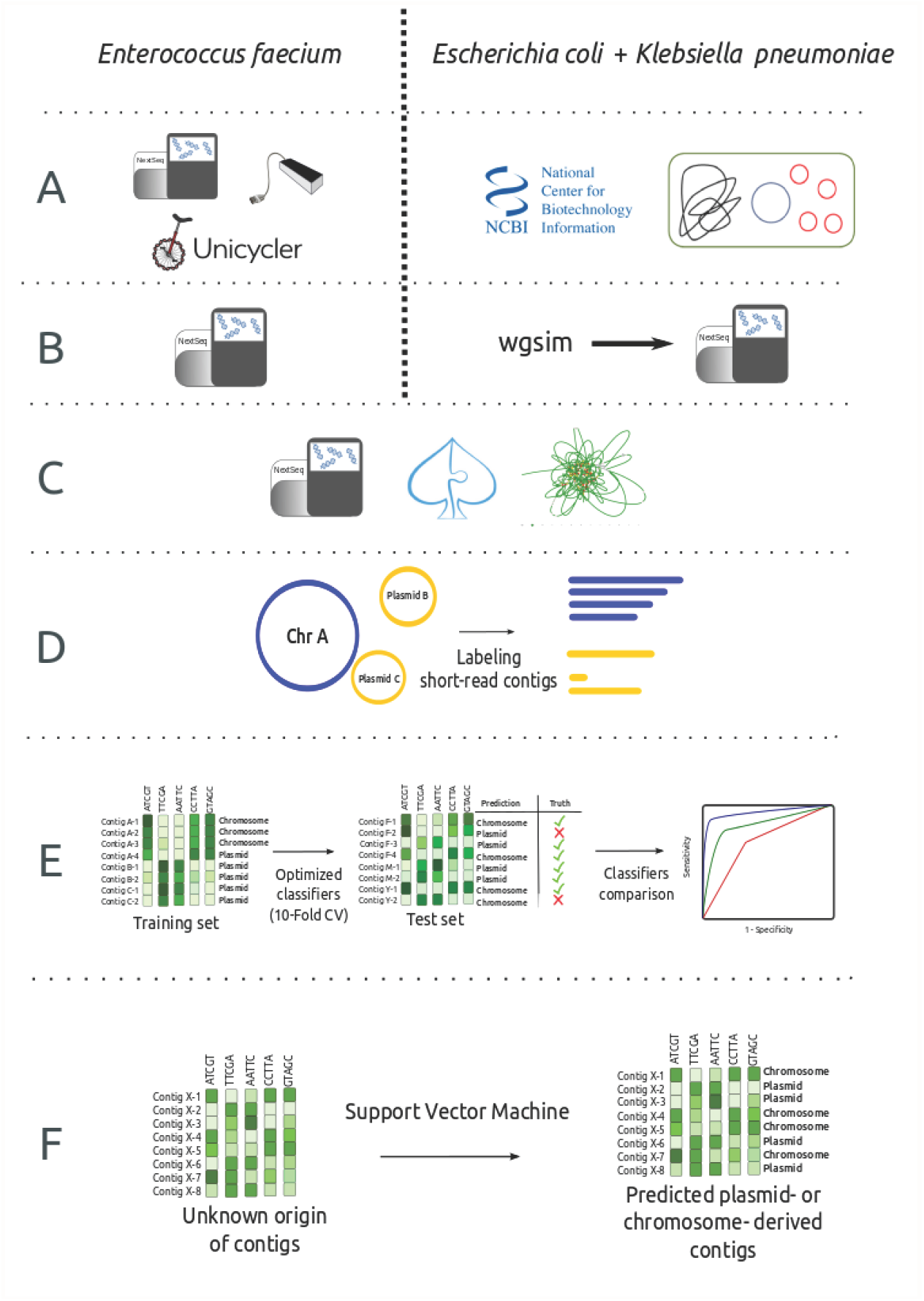
Workflow to create the plasmid models for *Enterococcus faecium, Klebsiella pneumoniae* and *Escherichia coli*. A) For *E. faecium*, 60 Illumina sequenced strains were selected for ONT sequencing and Unicycler was used to extend the number of complete genomes available for this species. For *E. coli and K. pneumoniae*, we downloaded complete genomes with plasmids associated from Assembly Entrez NCBI database. B) For *E. coli* and *K. pneumoniae*, we simulated reads with 50X coverage and no error rate using wgsim. C) Illumina simulated and non-simulated reads were *de novo* assembled using SPAdes. D) We mapped short-read contigs against complete genome sequences to define a reliable dataset of short-read contigs as plasmid- or chromosome-derived. E) For each bacterial species, five machine-learning classifiers were trained (10-fold cross-validation) and compared using a specific bacterial species training and test set. F) SVM models were implemented in mlplasmids and used to predict plasmid- and chromosome-derived sequences in isolates with only short-read WGS data available. The complete workflow is available from https://gitlab.com/sirarredondo/analysis_mlplasmids

### Implementing our optimized classifiers in mlplasmids

For each bacterial species, we selected the best model based on the resulting F1-score to predict plasmid- and chromosome- derived sequences. We implemented them in a new R package called ‘mlplasmids’ available at https://gitlab.com/sirarredondo/mlplasmids under the GNU General Public License v3.0. We also developed a Shiny app available at https://sarredondo.shinyapps.io/mlplasmids/ to enable plasmid-prediction with a graphical user interface (28).

### Comparison of mlplasmids against other plasmid prediction tools

We evaluated the performance of mlplasmids against PlasFlow (version 1.0), PlasmidSPAdes (version 3.8.2) and cBar (version 1.2). We considered contigs derived from isolates described in section ‘Defining an independent set of isolates’ to benchmark the algorithms. cBAR was run using default parameters. PlasFlow was run using a minimum posterior probability of 0.7 to assign a particular contig to a certain class and contigs labeled as ‘unclassified’ were not considered in this comparison. PlasmidSPAdes (version 3.8.2) generates its own assembly and resulting contigs were labeled as true- or false-positive results following the methodology described in section ‘Labeling short-read contigs as chromosome- or plasmid-derived’. For all the tools, we filtered out contigs with a length shorter than 1,000 bp.

We benchmarked above plasmid tools using: accuracy, f1-score, and precision (as previously defined in section ‘Building a machine-learning model’). PlasmidSPAdes does not predict chromosome-derived contigs thus we could not calculate its accuracy and F1-score. To overcome this, we used Quast (version 4.1) to map plasmid-predicted contigs against their respective complete plasmid sequences. We then retrieved the reported ‘genome fraction’ which consisted of the percentage of aligned bases in the reference genome to obtain an estimation of classifiers sensitivity (29).

### Validating mlplasmids against complete plasmid sequences

We used *K. pneumoniae* (n = 10) and *E. coli* (n = 3) complete genomes described in section ‘Defining an independent set of isolates’ to observe mlplasmids performance predicting complete chromosomal and plasmid sequences. Complete genomes of *E. faecium* from Assembly Entrez NCBI database (n = 24) were not included in the training set and were therefore considered to benchmark mlplasmids performance in complete *E. faecium* genomes.

### Predicting the location of antibiotic resistance genes

All assemblies of *E. faecium* (n = 369), *K. pneumoniae* (n = 1,346) and *E. coli* (n = 5,234) with an assembly level corresponding to ‘Contig’ were downloaded from NCBI Genomes FTP (ftp.ncbi.nlm.nih.gov/genomes/). For each downloaded draft assembly, we used Abricate (version 0.8.2) (https://github.com/tseemann/abricate) to screen contigs against the ResFinder database (release from 18th May of 2016) (30) to determine the presence of antimicrobial resistance genes. Abricate was run using a minimum DNA identity of 95% and a minimum coverage of 80%. To assign a particular contig as plasmid- or chromosome-derived, we used *E. faecium, K. pneumoniae* and *E. coli* SVM models in mlplasmids specifying a minimum posterior probability of 0.7 and a minimum contig length of 1,000 bp (Supplementary File 4).

We used isolates described in section ‘Defining an independent set of isolates’ to validate mlplasmids potential to predict the genomic context of a particular antibiotic resistance gene. We followed the same procedure as explained above to assign a resistance gene as plasmid- or chromosome-encoded. We used identical metrics introduced in section ‘Building a machine-learning model’ but considering genes as units.

### Predicting the plasmidome content of *E. faecium*

We used the *E. faecium* optimized model to predict plasmid- and chromosome-derived contigs from the collection of 1,644 *E. faecium* Illumina-sequenced (MiSeq/NextSeq) isolates. We filtered out contigs with a length shorter than 500 bp and labeled each contig as ‘plasmid’ or ‘chromosome’ derived based using the class with a highest posterior probability. SPAdes assembly statistics from this collection are shown in Supplementary Table S1.

## Results

To ensure that the new classifiers to predict plasmid- and chromosome-derived sequences were built using genome sequences from a large and diverse set of isolates belonging to each species, we first assessed the diversity present in our collections of *E. faecium, K. pneumoniae* and *E. coli*. We used Mash to sketch and cluster all the retrieved isolates from *E. faecium, K. pneumoniae* and *E. coli.* For *E. faecium,* we defined three main clusters and observed that our set of *E. faecium* (n = 60) expanded diversity present for complete genomes available in Assembly Entrez NCBI database (n = 24) (Suppl. Figure S2). Furthermore, seven isolates were part of a cluster in which we did not find NCBI complete genomes. Strikingly, we observed a single unique NCBI complete genome forming an independent cluster (GCF_000737555), corresponding to *Enterococcus faecium* T110, a probiotic strain (Suppl. Figure S2). For *K. pneumoniae*, we also observed and defined three main clusters from all complete genomes available in Assembly Entrez NCBI (n = 156). One of the three clusters was only composed by three *K. pneumoniae* isolates (GCA_000714635, GCF_000019565 and GCF_002156765) and showed a mash distance higher than 0.05 versus isolates present in the other two major clusters (Suppl. Figure S3). For *E. coli,* we observed three major clusters of isolates present in *E. coli* NCBI collection. As shown in Suppl. Figure S4 all defined *E. coli* clusters presented a high diversity versus each other in terms of mash distances. For *E. faecium* and *K. pneumoniae* the major clusters defined after hierarchical clustering showed lower diversity.

For each bacterial species, we used SPAdes contigs labeled as plasmid- or chromosome-derived and belonging to the test set to assess the performance of our optimized machine-learning classifiers. To investigate the applicability of genomic signatures as model features to distinguish between plasmid- and chromosome-derived sequences, we used Mash to sketch and cluster all the complete chromosomal and plasmid sequences of *E. faecium, K. pneumoniae* and *E. coli* that were included in this study. We observed that computed Mash distances using a k-mer size of 16 provided a clear separation between plasmid and chromosome sequences for each species (Figure 2 and Suppl. Figure S1). Additionally, we observed that some plasmid sequences from *E. coli* and *K. pneumoniae* were clustering together which suggested that plasmids from these two species share a high fraction of k-mers as a potential result of plasmid transmission between both species (Figure 2). This analysis confirmed that k-mer signatures can be used as classifier features to chromosome and plasmid sequences for single species. To develop a novel plasmid prediction tool, we used pentamer frequencies to train and test: Logistic regression, Bayesian classifier, Decision trees, Random forest and Support-Vector Machine classifiers since this resulted in a good ratio between number of objects and features (∼ 10 objects per feature). In addition, pentamer frequences were also previously used to distinguish plasmid sequences in metagenomic samples. Other potential plasmid features such as presence of known plasmid replication genes, contig coverage or contig length did not increase the performance of the algorithms and made the usability of the resulting classifiers less generalizable.

**Figure 2.**
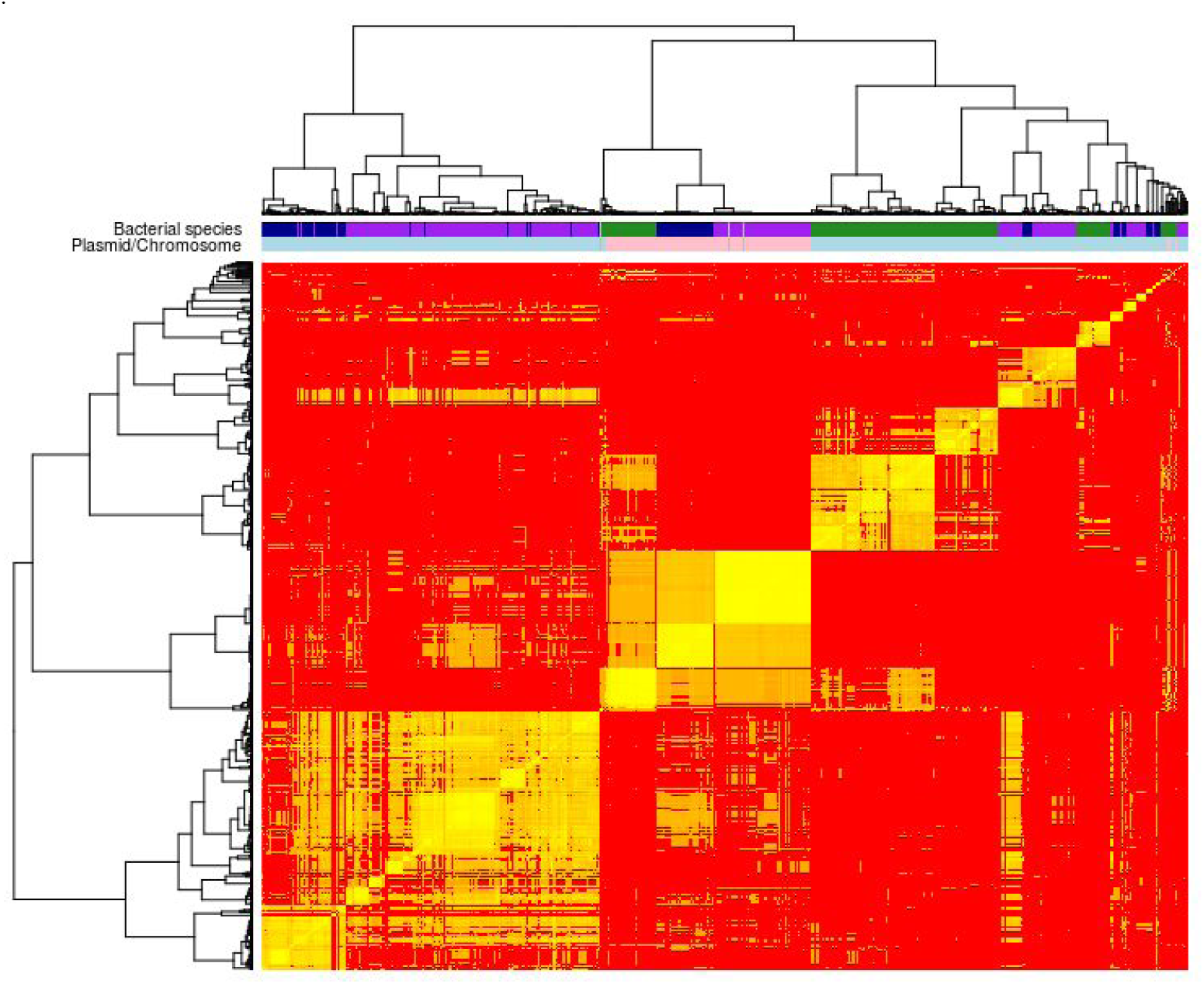
Ward hierarchical clustering of computed Mash distances (k = 16; s = 400) between all chromosome and plasmid sequences included in this study. Each node on the dendrogram corresponded to a either a plasmid (light blue) or chromosome (pink) sequence from *E. coli* (dark blue), *K. pneumoniae* (purple) or *E. faecium* (green).

Support-vector machine (SVM) was the machine-learning algorithm selected as best classifier for predicting plasmid-derived contigs in the three bacterial species. SVM performance in *E. faecium* (Accuracy = 0.94; F1-score = 0.92) and in *K. pneumoniae* (Accuracy = 0.92; F1-score = 0.90) was superior to the rest of machine-learning models and F1-score reflected that prediction of the model was balanced for both classes (Figure 3). In the case of *E. coli,* SVM performance (Accuracy = 0.95; F1-score = 0.76) reflected that prediction for the plasmid class was less accurate compared to the chromosome class (Sensitivity = 0.71) (Figure 3 and Supplementary File 3). This can be explained by a lower frequency of the plasmid class (Table 1) present in the training set of the machine-learning classifiers compared to the training sets of *Enterococcus faecium* and *Klebsiella pneumoniae* or a higher diversity from isolates categorized as belonging to this species (Suppl. Figure S4). For the three selected SVM models, we observed that metrics reported were higher when considering base-pairs as unit (Supplementary File 3). This indicated that misclassification mostly occurred on short length contigs (< 1 kbp) as shown for *E. faecium* SVM model (Suppl. Figure S5). We implemented *E. faecium, K. pneumoniae, E. coli* SVM models in a new R package called ‘mlplasmids’.

**Figure 3.**
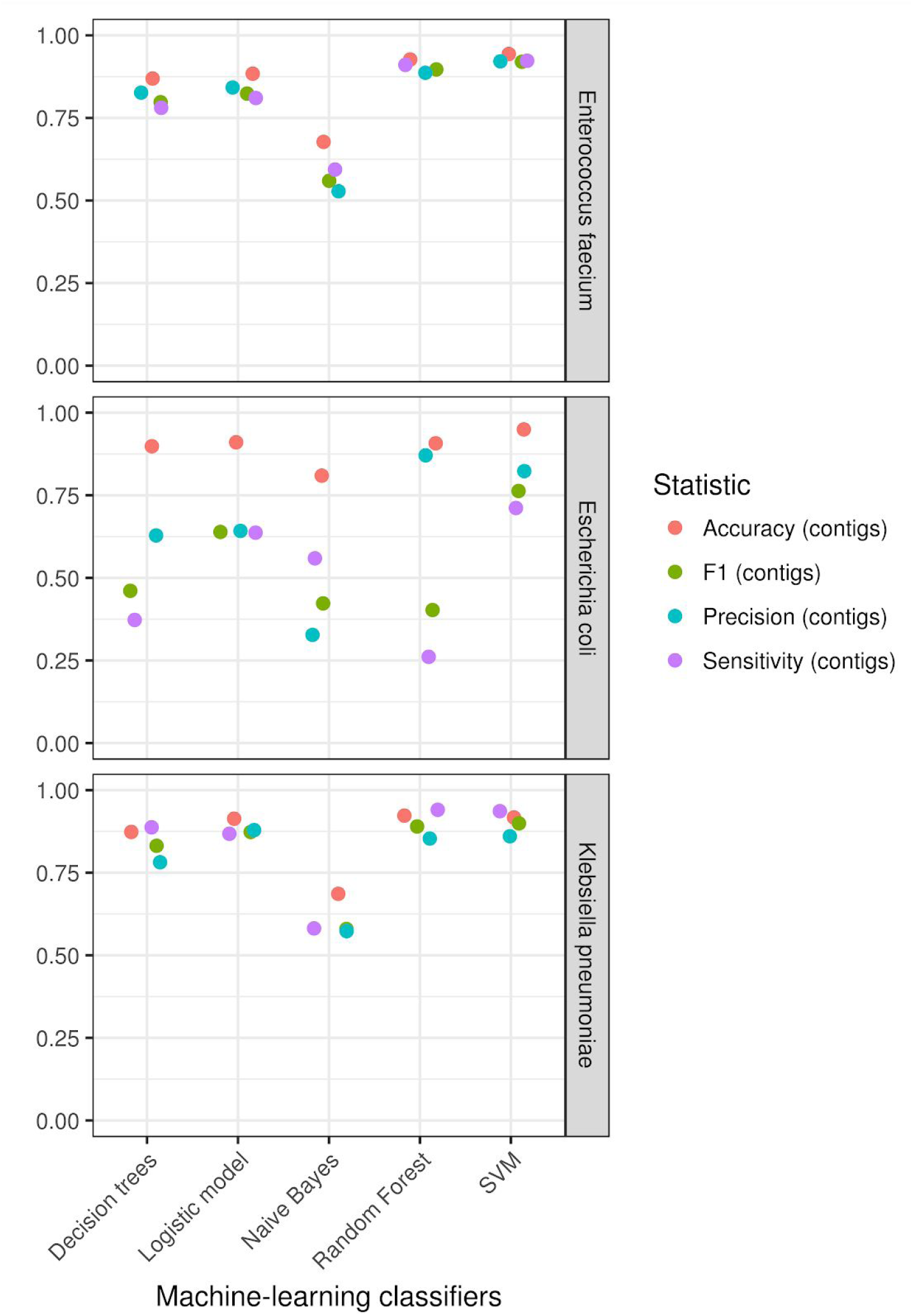
Performance of the optimized machine-learning classifiers: Decisiontrees, Logistic Model, Bayesian Classifier (Naive Bayes), Random Forest, and Support-Vector Machine(SVM) using our test sets for *E.faecium, E.coli* and *K.pneumoniae*. Statistics reported: accuracy(red), f1-score(green), precision(blue) and sensitivity (purple) are indicated using contigs as performance measure.

We benchmarked mlplasmids against other fully automated plasmid prediction tools using an independent set of isolates for which SPAdes contigs from non-simulated Illumina reads were available (*E. faecium* = 7, *K. pneumoniae* = 10, *E. coli* = 3). Performance of mlplasmids in *E. faecium* (F1-score = 0.94, Precision = 0.95) and *E. coli (*F1-score = 0.84, Precision = 0.88) was higher than cBAR, PlasFlow and PlasmidSPAdes (Figures 4 and 5). In the case of *K. pneumoniae*, mlplasmids metrics were overall better (F1-score = 0.88, Precision = 0.86) even though performance of plasflow (F1-score = 0.82, Precision = 0.72) and PlasmidSPAdes (Precision = 0.79) was also good and in the case of *K. pneumoniae* strain KPN555 performance was better compared to mlplasmids (Figure 5).

**Figure 4.**
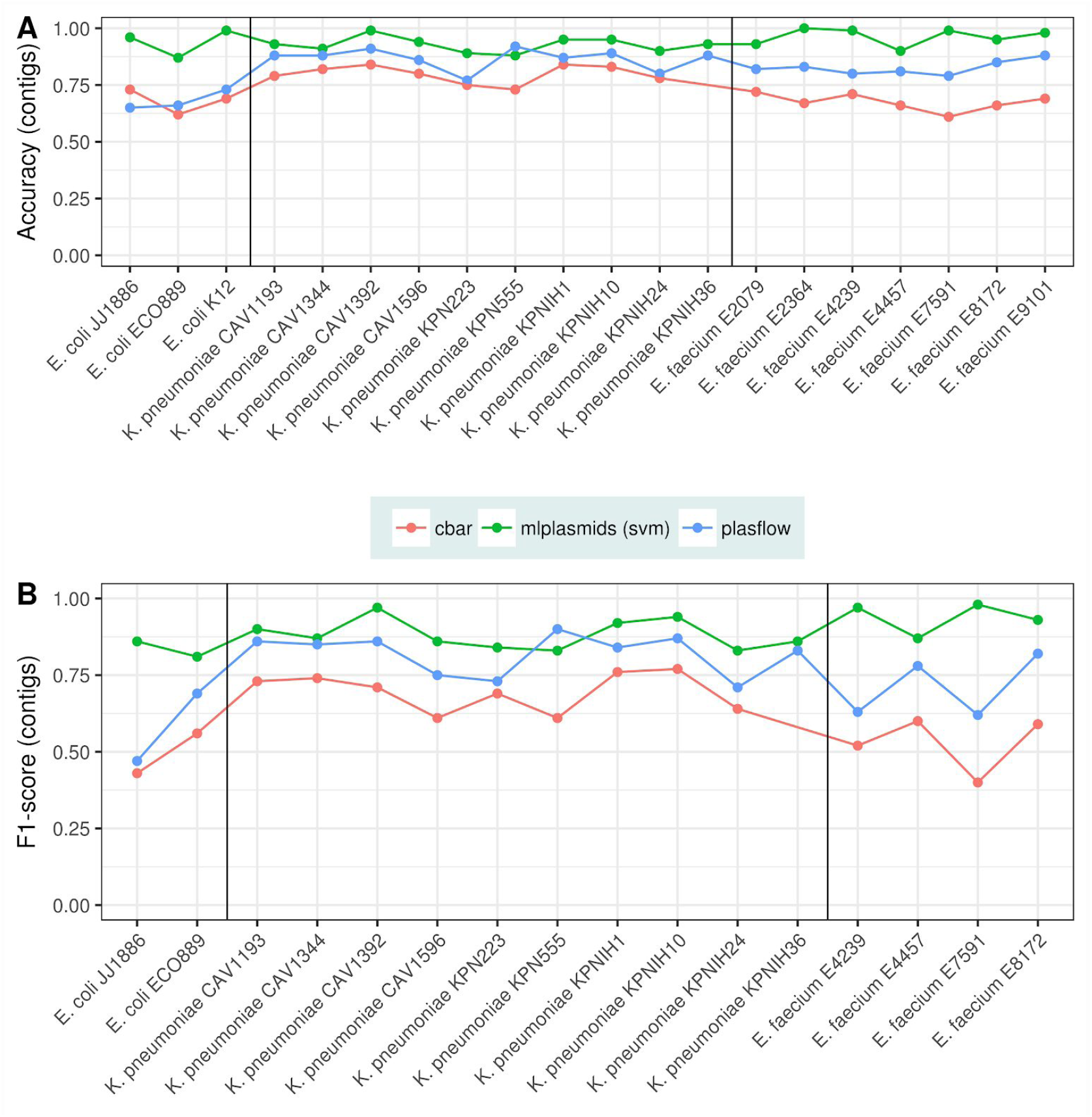
Benchmarking of cbar (red), mlplasmids (green) and plasflow (blue) using an independent set of isolates (n = 20). A) Accuracy was measured in contigs and reported for all isolates including samples considered as negative controls (*Escherichia coli* K. 12, *E. faecium* E2079, *E. faecium* E2364 and *E. faecium* E9101). B) F1-score was measured in contigs and only reported for isolates bearing plasmids (n = 16).

The average genome fraction values of mlplasmids for *E. faecium* (81.5%), *K. pneumoniae* (82.3) and *E. coli* (83.7%) indicated that most of the bases from the reference plasmids were covered in mlplasmids’ prediction even though contigs with a contig length smaller than 1,000 bp were filtered out (Figure 5B). For *K. pneumoniae,* overall genome fraction of plasflow (83.1) was higher than mlplasmids but precision (0. 72) indicated that a fraction of chromosomal-derived contigs was wrongly predicted as plasmid in the prediction (Figure 5A).

**Figure 5.**
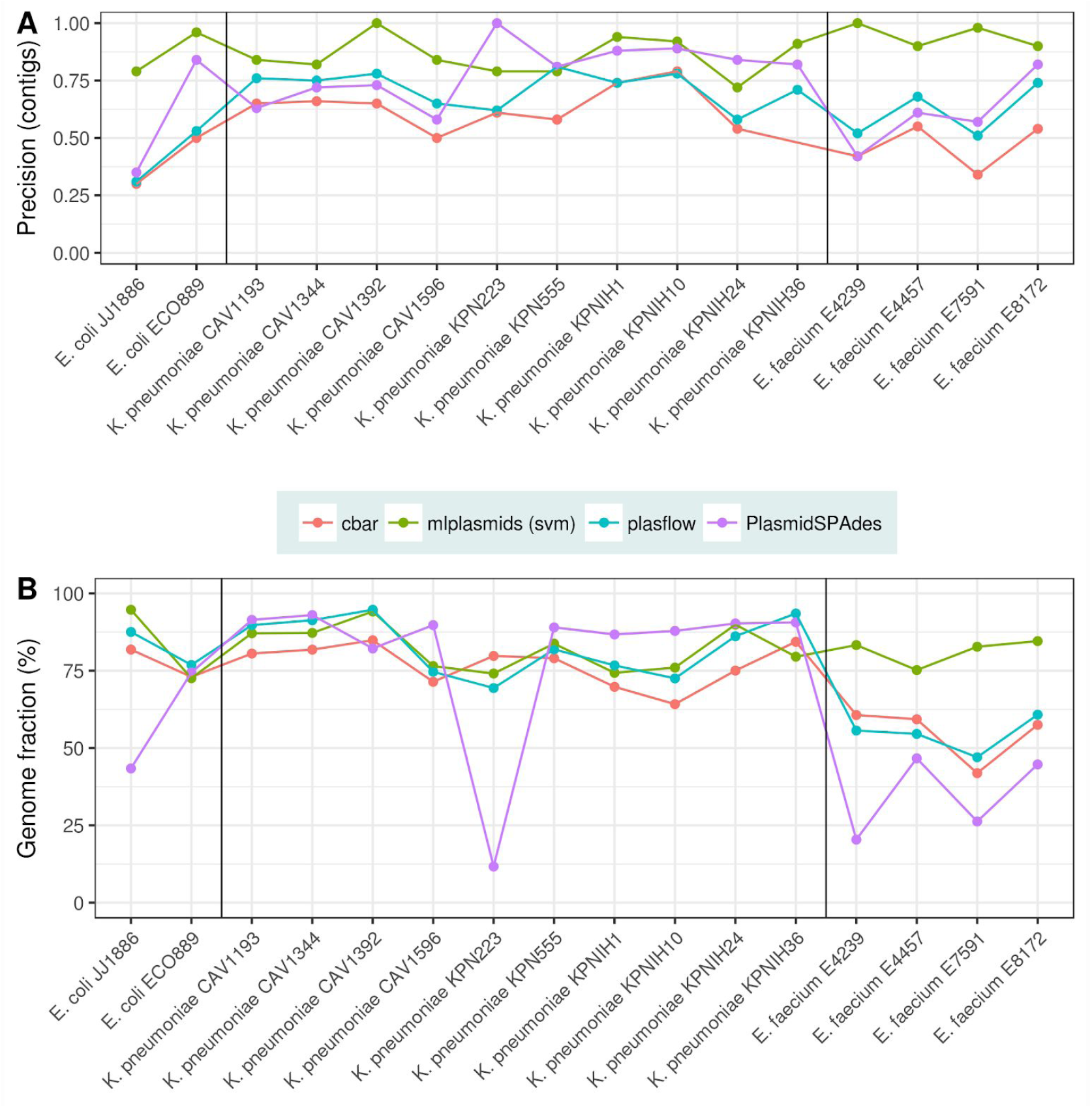
Comparison of cbar (red), mlplasmids (green), plasflow (blue) and PlasmidSPAdes (purple) using an independent set of isolates. A) Precision was measured in contigs and reported only for isolates bearing plasmids (n = 16). B) Genome fraction (measured as percentage of base-pairs) was extracted from Quast analysis for isolates bearing plasmids (n = 16).

Our approach of training the classifiers on datasets from single species was fundamental to obtain a good precision. This was also reflected in mlplasmids prediction for isolates corresponding to negative controls. For *E. coli* strain K-12 only a single contig (> 1000 bp) was erroneously predicted as plasmid-derived. We also observed similar very low numbers of false-positive plasmid assigned contigs for *E. faecium* E2079 (n = 6), *E. faecium* E9101 (n = 1) and for *E. faecium* E2364 (n = 0) all chromosome-derived contigs were predicted correctly.

Additionally, we evaluated mlplasmids using complete genome sequences from *E. faecium* (chromosomes = 24; plasmids = 82), *K. pneumoniae* (chromosomes = 10; plasmids = 33) and *E. coli* (chromosomes = 3; plasmids = 7) as input. Observed mlplasmids performance for *E. faecium* (F1-score = 0.99), *K. pneumoniae* (F1-score = 0.98) and *E. coli* (F1-score = 0.92) suggested that mlplasmids can also be used to predict plasmid and chromosomal sequences with a size larger than the average contig length obtained by WGS short-read data assembly. For all bacterial species, mlplasmids did not recover any false positive sequences (Specificity = 1). For *E. coli*, only a single plasmid sequence (NC_022662.1) was wrongly predicted as chromosome-derived but with a low posterior probability associated to that class (0.53). In the case of *K. pneumoniae*, mlplasmids misclassified a plasmid sequence with a length of 26.45 kbp (NZ_CP015133.1) from *K. pneumoniae* strain KPN555. For *E. faecium* two sequences were misclassified as chromosomal (NZ_LT598665.1 and NZ_CP019991.1) and of these sequences (NZ_CP019991.1) could correspond to a phage since its NCBI annotation showed two phage-related genes. This demonstrated the flexibility of mlplasmids to predict sequences from different lengths and may facilitate the classification of contigs generated from incomplete hybrid or long-read assemblies as exemplified for isolate *E. faecium* E7070. This isolate was selected for ONT sequencing and after hybrid assembly, 16 contigs were reported. Contigs predicted as plasmid by mlplasmids (n = 6) contained circularization signatures whereas the rest of the contigs (n = 10) were predicted as chromosome-derived (Suppl. Figure S6). This aided in designing appropriate PCR reactions to complete the genome sequence for E7070.

To show the potential of mlplasmids in determining if a particular gene of interest is plasmid- or chromosome-encoded, we predicted the location of antibiotic resistance genes in *E. faecium, K. pneumoniae* and *E. coli*. Firstly, we determined resistance genes in NCBI draft assemblies by using Abricate to screen contigs against the ResFinder database. Secondly, we used *E. faecium, K. pneumoniae* and *E. coli* SVM models in mlplasmids to determine whether these resistance genes were located in plasmid- or chromosome-derived contigs. For each identified resistance gene, we calculated the frequency of finding that particular gene on a predicted plasmid- or chromosome-derived contig.

For *E. faecium*, we assigned a total of 1,058 and 1,836 genes as chromosome- and plasmid-located respectively. We observed that most aminoglycoside resistance genes (e.g. *ant(6)-Ia_2*) were mainly present in a plasmid context (Suppl. Figure S7). Erythromycin resistance genes were preferentially present in one genomic context depending on the gene variant as exemplified by *erm(A)_1* and *erm(B)*_18 (Figure1). As previously described, the *vanA* operons were only present in plasmid-predicted contigs (31). Furthermore, *vanB* operons were present in both plasmid and chromosomal contexts but the frequency of chromosome-derived contigs was higher (∼ 0.73) (Suppl. Figure S7) (32). Validation of the prediction on *E. faecium* independent isolates (n = 7) revealed that all resistance genes (n = 43) predicted by Abricate were correctly predicted either as plasmid- or chromosome-derived (F1-score = 1.0).

For *K. pneumoniae*, we assigned a total of 5,107 and 10,432 ResFinder hits as chromosome- and plasmid-located. Most of antibiotic resistance genes showed a clear tendency of being mostly present in either a plasmid or chromosomal genomic context (Suppl. Figure S8). As described before (33), we observed some notable exceptions such as *armA* or *blaCTX-M-14_1* in which these particular resistance genes were also present in predicted chromosome-derived contigs (Suppl. Figure S9A). We performed the same analysis on *K. pneumoniae* isolates belonging to the independent set (n = 10) resulting in a total of 41 and 75 genes predicted as plasmid- and chromosome-encoded. Mlplasmids evaluation revealed that all predicted plasmid-encoded genes were correctly assigned (Precision = 1.0) and only five genes were misclassified as chromosome-encoded (F1-score = 0.96, Sensitivity = 0.93).

For *E. coli*, we assigned a total 4,517 and 8,085 ResFinder hits as chromosome- and plasmid-located. In contrast to *K. pneumoniae*, we observed that resistance genes were frequently identified in both plasmid and chromosomal contexts (Suppl. Figure S10). We also observed differences in gene location between resistance gene variants as exemplified for *qnrS2_1* which was frequently encoded in predicted plasmid-derived contigs in contrast to *qnrS1_1* which can be found in both genomic contexts (Suppl. Figure S9B). Interestingly, *mcr-1_1* was found in both plasmid and chromosomal contexts in *E. coli,* whereas for *K. pneumoniae* this resistance gene was only identified in plasmid-derived contigs (Suppl. Figure S9). Chromosomal locations of mcr-1_1 for *E. coli* have been described before (34). We predicted a total of 15 resistance genes from *E. coli* isolates which belonged to the independent set (n = 3). Mlplasmids performance revealed that 4 genes which were encoded in a single contig from *E. coli* ECO889 were wrongly predicted as chromosome-encoded whereas gene assignment was flawlessfor *E. coli* JJ1886 (F1-score = 0.88, Sensitivity = 0.80). As observed for *E. faecium* and *K. pneumoniae*, all genes predicted as plasmid-encoded were correctly assigned (Precision = 1.0).

Finally, we demonstrate mlplasmids utility predicting the plasmidome content of *E. faecium*. We predicted plasmid-derived sequences from a collection of 1,644 Illumina sequenced *E. faecium* isolates (Supplementary Table S1). Mlplasmids prediction using our R package took 1624,509 seconds (∼ 27 minutes) on a Linux laptop (Ubuntu 14.04) using a single core. Classifier prediction resulted in 194,884 contigs originating from the chromosome and 94,485 contigs with a predicted plasmid-origin in 1,640 isolates. Mlplasmids did not predict any plasmid-derived contig in four strains including one of our negative controls (‘*E. faecium* isolate E2364’). This corresponded to an average number of ∼119 chromosome - and ∼58 plasmid-derived contigs per isolate. Average cumulative length of chromosome and plasmid predicted contigs was 2,632,470 bp and 254,700 bp respectively. The average posterior probability of the predicted chromosome-derived contigs corresponded to 0.95 versus an average posterior probability of 0.91 for plasmid-predicted contigs (Suppl. Figure S11). This suggests a high likelihood that contigs were correctly assigned to each class.

Mlplasmids was implemented as a new R package and a Shiny app was developed and integrated in a web-server (Figure 6). As mlplasmids models use pentamer frequencies to predict plasmid-derived contigs, users can concatenate genome assemblies from single species in a single file which can facilitate the analysis of large collections of isolates. Assemblies can be uploaded to our web-server as.tar.gz. Users must select the species model (*Enterococcus faecium, Klebsiella pneumoniae or Escherichia coli*) for the plasmid prediction. After uploading a genome assembly, results appear as a tabular data in which each row corresponds to a sequence present in the fasta assembly. Additionally, results can be filtered using three options: i) minimum sequence length to report prediction; ii) minimum posterior probability for assignation of plasmid class; iii) minimum posterior probability for assignation of chromosome class. Results can be downloaded as.csv/xslx format.

**Figure 6.**
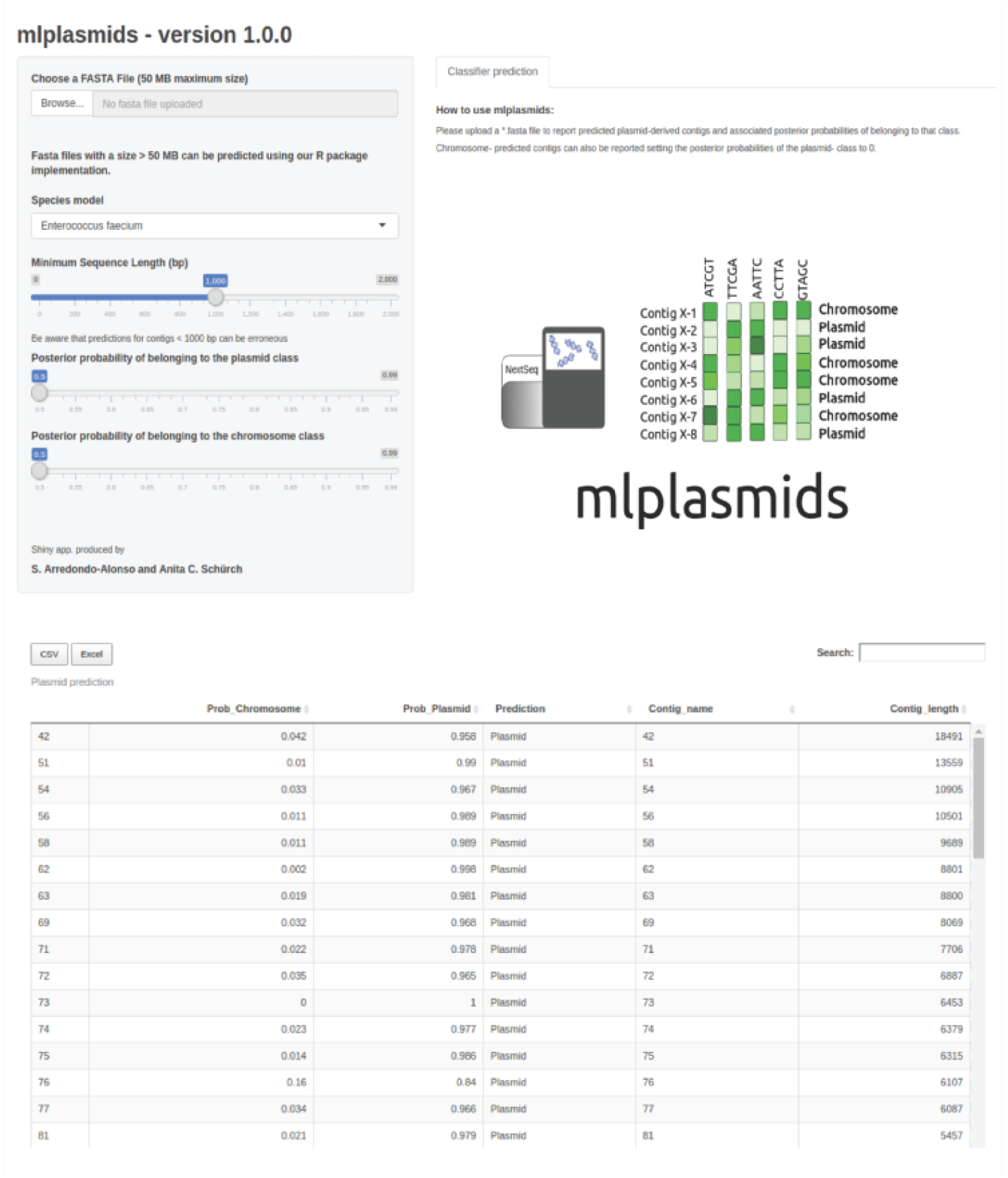
Graphical interface of mlplasmids web-server.

## Discussion

Here, we present a set of species-specific machine-learning classifiers to classify plasmid-derived contigs for three clinically relevant species: the Gram-positive bacterium *E. faecium,* and the Gram-negative bacteria *K. pneumoniae* and *E. coli.* We used genomic structure information from complete genomes to label short-read contigs as plasmid- or chromosome-derived and used them to train and test five different popular machine-learning algorithms. Using exclusively k-mer frequencies as input, we were able to predict whether short-read contigs from these three different species were plasmid- or chromosome-derived. Genome signatures were previously used in cBAR and more recently for PlasFlow to predict plasmid sequences from primarily metagenomes (12, 15). However, we show that mlplasmids performance is superior to cBAR and PlasFlow when predicting genome assemblies from *E. faecium, K. pneumoniae* or *E. coli*. This suggests that prediction accuracy increases substantially when training the classifiers with sequences from a single species. We additionally showed the importance of training our machine-learning classifiers using datasets that capture most of the species diversity. This was exemplified for *E. faecium* in which our long-read sequencing selection allowed to increase diversity and number of complete genomes resulting in a superior performance of our resulting model compared to current plasmid tools relying on machine learning. Mlplasmids however also outperformed PlasmidSPAdes, which relies on differences in coverage between plasmids and chromosome in the prediction of plasmid-derived contigs.

Our model can be used as a basis for plasmid classification for other tools such as PlacnetW, facilitating the reconstruction of plasmid sequences in a network graph, or for PlasmidSPAdes by filtering of chromosome-derived contigs regardless of contig coverage. We showed that the performance of mlplasmids was independent of sequence size which can facilitate the assignment of contigs derived from incomplete hybrid or long-read assemblies. We demonstrated the scalability of our model by predicting the plasmidome of a large collection of Illumina sequenced *E. faecium* isolates with high certainty. Furthermore, the low number of false-positive contigs predicted in isolates considered as negative controls suggests that the assignment of a gene of interest as plasmid- or chromosome-derived is possible regardless the presence of other plasmid signatures such as presence of plasmid replication genes, circularization signatures or contig coverage differences. We showcased a relevant application of mlplasmids, namely the assignment of resistance genes to an origin, plasmid or chromosome. Our prediction shows a high accuracy for all three species, and is, at least for *K. pneumoniae*, in accordance with previous reports (33).

To facilitate the usability of mlplasmids, we developed a Shiny app in which users can easily upload and retrieve mlplasmids prediction of genome assemblies. We anticipate that a similar methodology can be implemented to create new models for predicting plasmid- and chromosome-derived contigs for other bacterial species with a sufficient number of diverse and complete genomes.

